# DUSP5 Downregulation in Nucleus Accumbens Core Correlates with Synaptic Plasticity and Cue-Induced Cocaine Reinstatement

**DOI:** 10.1101/2025.09.19.677422

**Authors:** Juan Pablo Taborda-Bejarano, Michael Meyerink, Debbie C. Crans, Ramani Ramchandran, Constanza Garcia-Keller

**Author notes:** indicates equal contribution. Corresponding Authors: *Emails:* (neurobiological context of DUSP5 function), mailto (DUSP5 biology and ERK-related queries), (chemical biology and phosphatase). (CGK); (RR); (DCC).

## Abstract

The United States is currently facing a drug overdose epidemic, with substance use disorder (SUD) characterized by cyclical phases of drug use, withdrawal, and relapse. The nucleus accumbens core (NAcore), a brain region critical for reward and aversion behaviors, undergoes structural and functional synaptic adaptations in response to chronic drug exposure. These changes, particularly in dopamine D1 receptor-expressing medium spiny neurons (D1-MSNs), are implicated in drug-seeking behaviors and synaptic plasticity. However, the molecular mechanisms underlying these adaptations remain poorly understood. In this study, we investigate the role of dual-specificity phosphatase 5 (DUSP5), an phosphatase known to deactivate extracellular signal-regulated kinase (ERK), in cocaine-induced neuroplasticity. While prior research has linked other DUSP family members to various drugs of abuse, the specific role of DUSP5 in cocaine addiction remains unexplored. We hypothesized that lack of DUSP5 contributes to NAcore synaptic plasticity during cue-induced cocaine reinstatement. To test this, we employed a rat cocaine self-administration model, integrated molecular analyses, and mined publicly available single-cell RNA sequencing data from cocaine-treated NAcore. Our findings aim to elucidate the potential involvement of DUSP5 in cocaine-related synaptic adaptations and behavior, addressing a significant gap in the mechanistic understanding of SUD.

## INTRODUCTION

Cocaine addiction continues to pose a major public health challenge in the United States. In 2021, an estimated 1.4 million individuals met the criteria for cocaine use disorder. Alarmingly, the rate of cocaine-related overdoses is increasing, largely due to the combined use of cocaine with synthetic opioids like fentanyl. This pattern of polysubstance abuse marks the so-called “fourth wave” of the opioid crisis, defined by the convergence of stimulant and opioid use [1, 2]. The rising prevalence of stimulant use has introduced additional complications, including adverse behavioral outcomes such as neurological impairments, suicidal ideation, psychosis, aggression, and violence, while also complicating treatment approaches [1–3].

Clinical evidence underscores the need for a deeper understanding of how drugs alter brain function to effectively treat substance use disorder (SUD) [4]. SUD is closely linked to synaptic changes within the nucleus accumbens (NAc), a brain region central to reward processing, motivation, and addiction [5]. The NAc, or ventral striatum, contains core and shell subregions that play distinct roles in addiction-related behaviors [6]. The core is primarily associated with driving goal-directed actions and drug-seeking (reinstatement) behavior whereas the shell is involved in motivational processes and the formation of stimulus-reward associations [6, 7]. Within the NAcore, approximately 90% of neurons are medium spiny neurons (MSNs) that express either D1 or D2 dopamine receptors [8]. Activation of D1-MSNs has been shown to facilitate motivated behaviors [9], whereas D2-MSN activation tends to suppress them. Long-lasting, experience-dependent changes in synaptic strength within the NAcore are believed to underlie drug-induced neuroplasticity [7, 10]. This plasticity is evident both structurally, through changes in dendritic morphology, and functionally, via altered glutamatergic receptor signaling, distinct measures that nonetheless influence one another [11, 12]. Despite significant efforts, therapeutic strategies aimed at reversing drug-induced synaptic plasticity have yielded limited success. This is due in part to an incomplete understanding of the mechanisms governing synaptic plasticity under both normal and drug-altered conditions. In the present study, we address this gap by investigating a novel molecular target: the dual-specificity phosphatases (DUSPs), with a particular focus on DUSP5, a member of this family, and its role in SUD.

DUSPs are dual specificity phosphatases in that they remove phosphate residues (dephosphorylation) from serine (S), threonine (T) and tyrosine (Y) residues. Indeed, DUSP5 removes phosphate groups (pT and pY) from extracellular regulated kinase (ERK) to inactivate the kinase [13–15]. *DUSP5* and *DUSP6* mRNA expression is upregulated (~1.4-fold) in the prefrontal cortex following heroin abstinence[16]. Conversely, *DUSP5* expression in the NAc decreases (~0.67-fold) shortly after heroin injection [17]. Methamphetamine exposure also affects DUSP expression: *DUSP1* and *DUSP6* mRNAs and protein levels were increased in multiple brain regions, including the cortex, striatum, thalamus, and hippocampus right after a single injection [18, 19]. Other DUSP family member implicated in SUD is DUSP15. *DUSP15* mRNA and protein expression was downregulated in NAc after 24 hs and 10 days after the last morphine injection [20]. DUSP15 over-expression using virus prevented the morphine-paired contextual memory, facilitated extinction and inhibited reinstatement and abolished ERK activation [20]. Additionally, a variant in the DUSP27 gene (rs950302) is associated with heroin addiction in African Americans [21]. Collectively, these findings suggest that DUSP genes may contribute to both vulnerability to and progression of addictive behaviors, potentially playing key roles in relapse by influencing drug-seeking behavior of psychostimulants and opioids.

Our team has been investigating DUSP5 in the vascular context and identified a novel molecular mechanism in its regulation of p-ERK [22–24]. Previous study [25, 26] have shown that DUSP5 is essential for regulating the hippocampal dentate gyrus neuroplasticity, a brain region important for learning and memory. Based on these findings, we hypothesized that DUSP5 may contribute to maladptive synaptic plasticity in the NAcore. To test this hypothesis, we employed a multidisciplinary approach combining molecular and behavioral methods. These included mining single-cell RNA sequencing data from NAcore tissue of cocaine-treated rats, conducting standard rat cocaine self-administration paradigms, analyzing dendritic spine morphology, and performing correlation analyses between behavior and structural synaptic plasticity.

## MATERIALS AND METHODS

### Animal housing and surgery

Nine male and female Long–Evans rats were individually housed using a 12/12 h light/dark cycle with ad libitum food and water in a temperature and humidity-controlled environment. All experimentation occurred during the dark phase, and animals were allowed to acclimate to the vivarium environment for a week before surgery. Rats were ~70 days old when they were anesthetized with isofluorane before implanting indwelling jugular catheters (surgical details have been described previously [27, 28], and received ketorolac as a preoperative analgesic. All procedures were in accordance with the National Institutes of Health Guide for the Care and Use of Laboratory Animals and the Assessment and Accreditation of Laboratory Animal Care. Our animal protocol at Medical College of Wiscon is ID: AUA00007874.

### Drugs and reagents used

Cocaine hydrochloride was supplied by the National Institute of Drug Abuse. NAcore MSNs were labeled with a virus (0.75 µl/hemisphere at a rate of 0.15 ml/min) that drove expression of membrane targeted mCherry under control of the synapsin promoter (pAAV2-shRNA, titer 1.39 × 10^15^ GC/ml). The AAV-shRNA vector consisted of a CMV promoter driving mCherry with an SV40 polyadenylation signal followed downstream by a U6 polymerase III promoter and polymerase III termination signal [29]. Rabbit anti-DUSP5 monoclonal antibody (Abcam Cat # ab200708, 1:500) and amplification of the viral reporter was performed at the same time using anti-mCherry (LSBio 204825, 1: 500). Secondary antibodies Alexa Fluor-conjugated to the appropriate species was used as well (Life Technologies, 1:1000).

### Self-administration (SA), extinction, and reinstatement

All procedures occurred in standard operant chambers equipped with two retractable levers, a house light, cue light, and 2900-Hz tone generator (Med Associates). Five male cocaine-treated and 4 male yoked saline rats were used in this study. Before cocaine SA training, animals were food deprived for 48 h, and then underwent a single 2-h food training session in which presses on the active lever resulted in the delivery of a single food pellet (45 mg, Noyes) on a fixed-ratio 1 (FR1) schedule of reinforcement. Following food training, animals were left with food *ad libitum* for the remainder of the experiments. One day later, animals began 2-h sessions cocaine SA on an FR1 schedule with a 20-s time out. Each active lever press resulted in an infusion of cocaine hydrochloride (0.25 mg/infusion) and simultaneously triggered a compound cue, consisting of a light positioned above the active lever and a 2900 Hz tone,serving as a conditioning stimulus. An inactive lever was also provided to control for non-motivated responding. Active lever presses made during the time out were counted but did not result in drug delivery. Rats underwent SA for a minimum of 10 days, until they met maintenance criteria of ≥10 infusions of cocaine over 10 days, as well as discrimination between active and inactive levers (>75% presses on active lever).

Following successful acquisition and maintenance of cocaine SA, extinction training (2 h/d) began. During extinction, presses on the previously active lever were recorded but no longer produced drug or presentation of the drug-paired cues. All rats underwent at least 8 days of extinction, or until two consecutive days revealed ≤25 active lever presses.

Cue-induced drug reinstatement training was conducted following extinction. During reinstatement sessions, animals were tested under a FR1 schedule, in which each active lever press resulted in presentation of the previously drug-paired light/tone cues used during SA, but without cocaine delivery. To initiate the session and signal cue availability, a single non-contingent (“free”) cue presentation was delivered at the start of the session. Thereafter, cue presentations occurred only in response to active lever presses. For dendritic spine analysis, both saline- and cocaine-exposed animals were removed from the operant chambers and euthanized 15 minutes after the start of the reinstatement session. A subset of animals served as yoked-saline controls and received a noncontingent saline infusion paired with light and tone cues according to a pre-programmed pattern of responding based on the cocaine average, but were otherwise treated identically throughout. All behavioral sessions lasted 2 hours and were performed at the same time each day.

### Quantification of dendritic spine morphology and immunohistochemistry

Rats were anesthetized with isofluorane as per our approved animal protocol. Transcardial perfusions with PBS followed by 4% paraformaldehyde (PFA) in PBS. Brains were removed and placed in sucrose buffer for 24 hs, then coronally sectioned at 100 μm in PBS on a vibratome. For immunohistochemical detection of DUSP5, sections were incubated with a rabbit recombinant primary antibodies against DUSP5 (1:500) for 48 h at 4°C with gentle agitation (as described in [30]). mCherry viral reporter signal was amplified using mcherry antibody. Following incubation with primary antibodies, sections were washed and followed by overnight at 4°C secondary incubation with the appropriate species-specific Alexa Fluor-conjugated secondary antibody (1:1000, Life Technologies). After a brief PBS wash, tissue was mounted onto slides in aqueous medium mounting with ProLong gold Antifade (Life Technologies) to preserve fluorescence intensity over time.

Spine morphology was quantified as described in detail previously [31]. Briefly, images of MSN-labeled segments were taken on a confocal microscope (Leica SP8) using a 561-nm laser line. Images of dendrites were taken through a 63× oil immersion objective with a numerical aperture of 1.4, using a 3.5× digital zoom. Images were deconvolved via LASX imaging software before analysis, and a 3-D perspective was rendered by the Surpass module of Imaris software package version 9.9 (Bitplane). Final data set voxel dimensions were of 0.049 μm in the *xy* plane and 0.3 μm in the *z*. The smallest quantifiable diameter spine head was set to 0.14 μm. Only spines on dendrites beginning >75 μm and ending <200 μm distal to the soma and after the first branch point were quantified on cells localized to the NAcore. The length of quantified segments was 45–55 μm. DUSP5 protein immunoreactivity was quantified in the same dendritic segments used for morphological analyses. To assess colocalization of DUSP5 with virally labeled dendrites, the total volume (µm^3^) of DUSP5 puncta signal was measured and normalized to the volume (µm^3^) of the corresponding dendritic shaft or spine, yielding an index of DUSP5 signal intensity within the isolated dendritic compartments (see **Supplemental Fig. 1** for details). All imaging and subsequent analyses were conducted blinded to experimental group assignments.

### Statistics

All statistical analyses were performed using GraphPad Prism version 10. All measurements were conducted by experimenters blinded to treatment conditions. For drug SA, acquisition was defined as achieving a criterion of ≥10 infusions per 120-minute session across two consecutive days. Statistical analysis for SA data was conducted using two-way ANOVA with repeated measures over time. Cue-induced reinstatement was also analyzed using two-way ANOVA. Dendritic spine morphology was analyzed using Imaris (Bitplane), including measurements of medium spiny neuron (MSN) dendritic spine density, spine head diameter, and DUSP5 immunoreactivity in both the spine and shaft compartments. For dendritic spine and DUSP5 immunoreactivity analyses, data normality was assessed using the D’Agostino & Pearson normality test. Comparisons between groups were made using unpaired t-tests. Correlation analyses and linear regressions were performed using GraphPad Prism.

## RESULTS

### *DUSPs* mRNA levels changes in D1-MSN in NAcore isolated from non-contigent saline- and cocaine-injected rats

To examine how cocaine exposure affects DUSP family gene expression across different cell types in the NAc, we analyzed a publicly available single-nucleus RNA sequencing (snRNA-seq) dataset from rats treated with acute (1 injection) and repeated (7 injections) intraperitoneal cocaine [32]. NAc tissue was collected 1 hour after the final injection, and nuclei were isolated for transcriptomic profiling (**Fig. 1A**). Data from acute and repeated treatment groups were integrated to examine cocaine-induced changes in mRNA expression across 16 annotated brain cell populations, including D1- and D2-expressing MSNs, interneurons, astrocytes, oligodendrocytes, and microglia. Among the DUSP family members examined (Dusp5, Dusp6, Dusp7, Dusp14, Dusp15, and Dusp27), only Dusp5 showed a selective increase in expression in D1-MSNs within the NAcore (Drd1-MSN-1), but not in NAc shell (Drd1-MSN-2), following cocaine treatment (**Fig 1B; blue box**). The difference between Drd1-MSN-1 and Drd1-MSN-2, as described in [32] lies in the varying expression levels of *Htr4* (which encodes the serotonin receptor 4) and *Calb1* (which encodes the calcium-binding protein Calbindin 1). These genes are enriched in the NAcore, indicating that the two Drd1-expressing MSN subtypes identified through snRNA-seq are distinct both transcriptionally and spatially. No changes in *Dusp5* expression were observed in D2-MSNs from either the NAc core or shell (Drd2-MSN-1 or Drd2-MSN-2), nor in other neuronal or glial cell types. Additionally, expression of the other DUSP genes remained largely unchanged across all cell populations analyzed. These results suggest that *Dusp5* mRNA is selectively upregulated in D1-MSNs of the NAcore following cocaine exposure, identifying DUSP5 as a potential cell type–specific regulator of cocaine-induced neuroadaptations.

**Figure 1.**
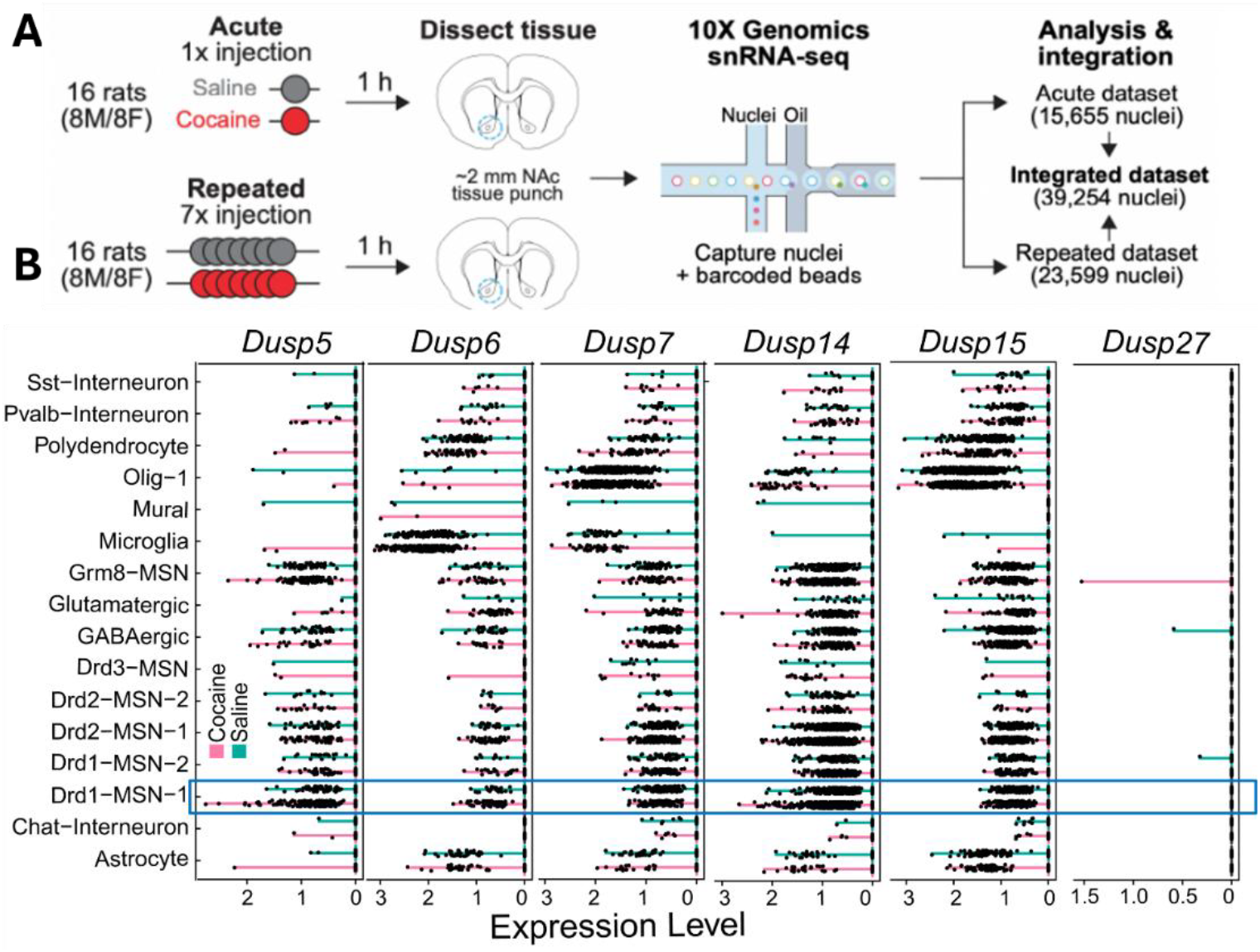
DUSP5 mRNA is selectively upregulated in D1-MSNs of the NAc following acute cocaine exposure. **Top:** Schematic of the experimental design from a publicly available single-nucleus RNA-sequencing (snRNA-seq) dataset [32]. Rats were treated with either acute (1x) or repeated (7x) intraperitoneal (i.p.) injections of cocaine or saline. One hour after the final injection, ~2 mm tissue punches from the NAc were collected for snRNA-seq using the 10x Genomics platform. The integrated dataset contains transcriptomic profiles from 39,254 nuclei, with data stratified by cell type. **Bottom:** Gene expression levels (log-transformed) for Dusp5, Dusp6, Dusp7, Dusp14, Dusp15, and Dusp27 across distinct brain cell populations, including Drd1- and Drd2-expressing MSN, interneurons, glial cells, and astrocytes. Pink lines represent expression following cocaine, and teal lines represent saline controls. Notably, Dusp5 mRNA is selectively upregulated in Drd1-MSNs (D1-MSNs) following acute cocaine administration, while other DUSPs show no consistent cocaine-related changes across cell types. This Dusp5-specific induction in D1-MSNs highlights its potential role in mediating early, cell type– specific transcriptional responses to cocaine exposure in the NAc.

### Cocaine-treatment decreased DUSP5 immunoreactivity in NAcore MSNs

We implanted intravenous catheter to allow SA of cocaine and microinjected AAV2-hSyn-hChR2-EYFP to label dendrites for later morphologic and immunoreactivity analysis of DUSP5 (**Fig. 2A**). Rats were trained to self-administer cocaine along with yoked saline controls (**Supplement Fig. 2A**). No statistical difference was observed between total number of saline infusions (flat rate) compared to cocaine-treated rats (**Supplement Fig. 2B**). After animals completed extinction training, reinstated cocaine seeking was induced for 15 min by restoring cocaine-paired cues (**Supplement Fig. 2C**). We and others have shown that this SA protocol induces synaptic plasticity in NAcore MSNs [10, 30, 31, 33], with some adaptations being specific to D1-MSNs [30, 34], providing the rationale for using this model in our study.

**Fig. 2.**
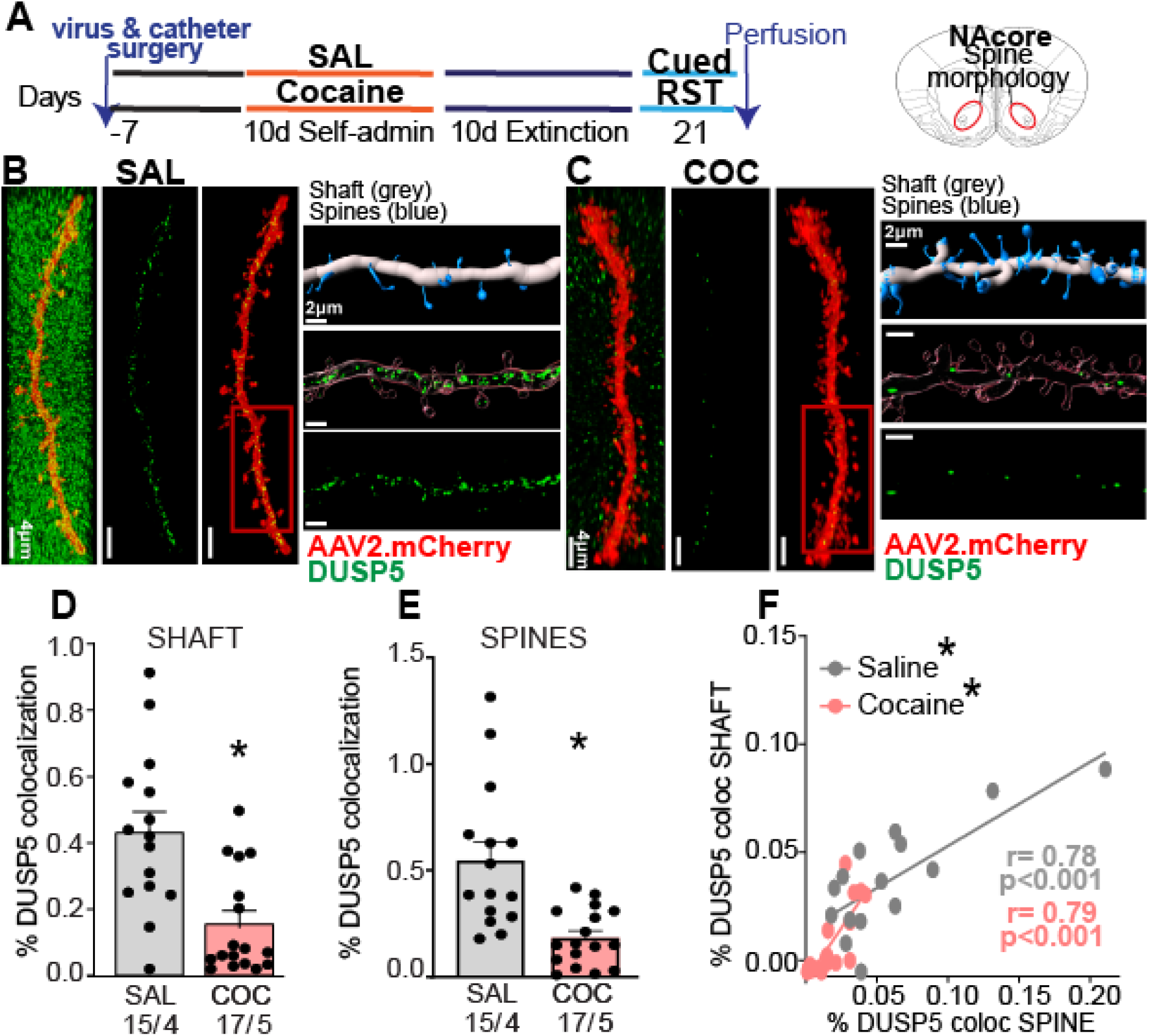
Reduced DUSP5 Protein Levels in NAcore MSNs During Cue-Evoked Cocaine Seeking. **A)** Experimental timeline illustrating the sequence of viral and catheter surgeries, SA training, and the time of perfusion for both cocaine (COC) and saline (SAL) groups. **B & C)** Representative dendritic segments from NAcore MSNs labeled with mCherry.synapsin (red) and DUSP5 (green) in SAL and COC groups. The three vertical panels show confocal images of DUSP5 puncta (green) and dendritic labeling (red): left—merged full image, middle—isolated dendrite, right—DUSP5 puncta localized within the dendrite. Scale bar = 4 µm. Next to it, horizontal panels display high-magnification insets of each segment. Also shown is the quantification method: top—3D Imaris Bitplane reconstruction distinguishing shaft (gray) and spines (blue); middle—isolated dendritic segment with DUSP5 puncta; bottom—quantification of DUSP5 colocalization in shaft and spines. **D)** Significant reduction of DUSP5 immunoreactivity in dendritic shafts of COC group compared to SAL group (unpaired t-test t_(30)_ = 3.902, *p* < 0.001). **E)** Significant reduction of DUSP5 immunoreactivity in dendritic spines of COC group compared to SAL group (unpaired t-test t_(30)_ = 4.032, *p* < 0.001). **F)** Positive correlation between DUSP5 immunoreactivity in dendritic shafts and spines in both SAL and COC groups (saline: r = 0.78, p < 0.001; cocaine: r = 0.79, p < 0.001). Data are shown as mean ± SEM. *N* shown as number of neurons quantified over number of animals in each condition; \**p* < 0.001 comparing COC to SAL.

Brain slices were prepared of NAcore and stained for DUSP5 (**Fig. 2B and C**). We quantified the DUSP5 immunostaining results using a 3D-rendering method previously published by our group [30] and described in (**Supplemental Fig. 1**). We found decreased DUSP5 immunoreactive puncta within NAcore dendrites after 15 min of cue-induced reinstatement in cocaine-treated animals, compared with levels of immunoreactive DUSP5 puncta from saline-treated rats. DUSP5 immunoreactivity was significantly reduced in shaft and spines after cocaine treatment (**Fig. 2D & E**). Furthermore, we observed a positive correlation between DUSP5 levels in shafts and spines across both treatment groups (**Fig. 2F**). These findings demonstrate that DUSP5 protein levels are diminished in both dendritic shafts and spines of NAcore MSNs during cue-induced cocaine reinstatement and suggest a coordinated regulation of DUSP5 immunoreactivity within these subcellular compartments.

### DUSP5 immunoreactivity in NAcore MSN correlates with active lever presses during reinstatement

To support a causal relationship between immunoreactivity of DUSP5 in shaft and spines and behavior we performed correlation analysis. Specifically, we examined association between DUSP5 inmunoreactive puncta and (1) total number of infusions, (2) infusions during the last five days of SA, and (3) active lever presses during cue-induced reinstatement. Simple linear regression revealed no significant correlations between DUSP5 immunoreactivity (in either spines or shafts) and total cocaine infusions (**Fig. 3A**), saline infusions (**Fig. 3D**), or infusions during the final five days of SA (**Fig. 3B and 3E**). However, a significant positive correlation was observed between DUSP5 immunoreactivity in dendritic spines and shaft with the number of active lever presses during reinstatement in cocaine-treated animals (**Fig. 3C**). No such correlation was found in saline-treated animals (**Fig. 3F**). These results suggest that DUSP5 immunoreactivity in NAcore MSNs is more closely associated with cocaine-seeking behavior during reinstatement than with the overall level of drug exposure, supporting a potential role for DUSP5 downregulation in mediating cue-induced relapse mechanisms.

**Fig. 3.**
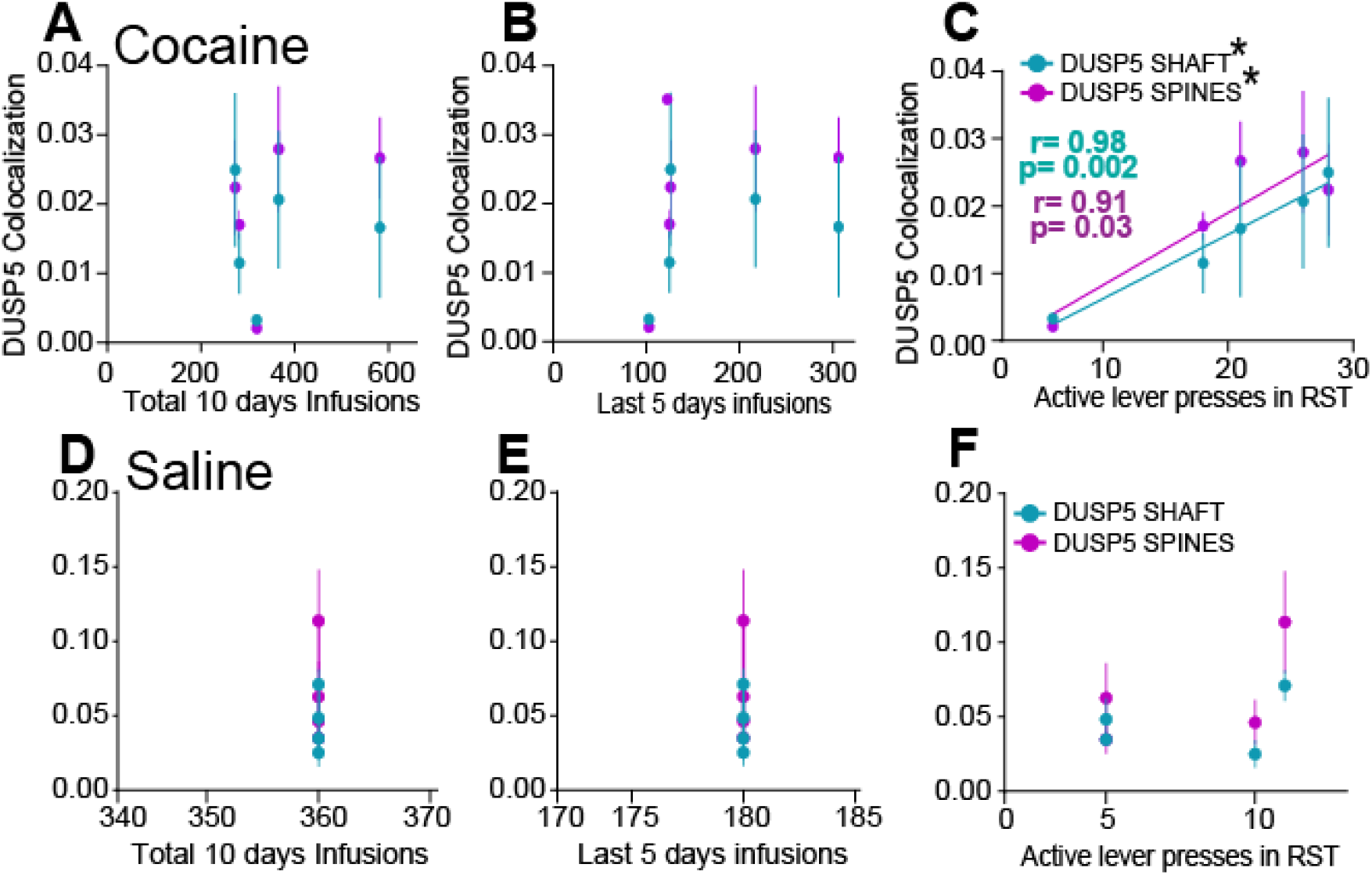
DUSP5 Immunoreactivity positively correlates with active lever pressing during cue-induced reinstatement. **A–B)** No significant correlation was observed between DUSP5 immunoreactivity in dendritic shafts or spines with **(A)** total cocaine infusions or **(B)** cocaine infusions during the last 5 days of SA. **C)** A significant positive correlation was found between DUSP5 immunoreactivity in spine heads and the number of lever presses during cue-induced reinstatement (spines: r = 0.91, p = 0.03; shaft: r = 0.98, p = 0.002). **D–E)** Replication of analyses in a saline cohort showed no correlation between DUSP5 immunoreactivity in shaft or spine heads and **(D)** total infusions or **(E)** infusions during the last 5 days. **F)** No correlation between DUSP5 levels in spine heads and lever pressing during reinstatement.

### Enhanced Synaptic Plasticity in NAcore MSNs During Cue-Induced Reinstatement

Previous work by our group and others have shown that cocaine drug-seeking induces structural and functional synaptic plasticity in NAcore MSNs[30, 35]. To assess this further, we analyzed dendritic spine morphology in the same dendritic segments used for DUSP5 immunoreactivity quantification (**Fig. 2**), providing a readout of synaptic plasticity (**Fig. 4**). We found a significant increase in spine density (spines per μm) in cocaine-treated animals compared to saline controls (**Fig. 4B**). In contrast, there was no significant difference in spine head diameter between cocaine- and saline-treated groups (**Fig. 4C**). Taken together, these results show that cocaine exposure enhances dendritic spine density, indicative of structural synaptic plasticity, in NAcore MSNs. Notably, these structural changes occurred in the same dendritic segments that exhibited reduced DUSP5 immunoreactivity, supporting an association between DUSP5 downregulation and maladaptive plasticity linked to cocaine-seeking behavior.

**Figure 4.**
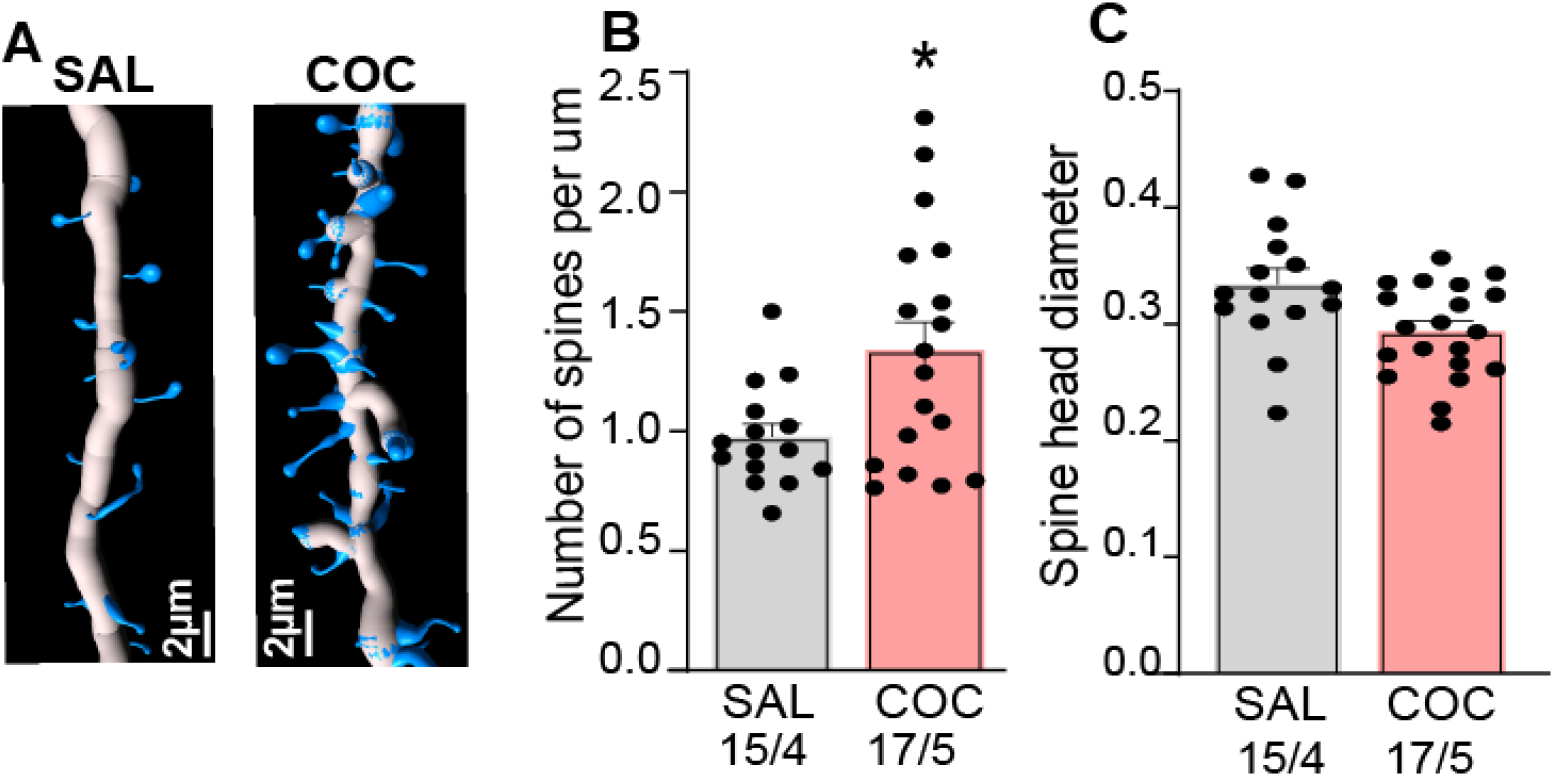
Enhanced synaptic plasticity in NAcore MSNs during cue-induced reinstatement is associated with reduced DUSP5 immunoreactivity. Synaptic plasticity was examined in the same animals analyzed in Figures 2 and 3. **A)** Representative dendritic segments from NAcore MSNs in saline- (SAL) and cocaine-treated (COC) rats. **B)** Increased number of spines per um (spines/µm) in cociane treated rats compared to saline group (unpaired t-test t_(31)_ = 2.99, *p* = 0.012). **C)** No significant difference in spine head diameter between saline- and cocaine-treated rats. Data are shown as mean ± SEM. *N* shown as number of neurons quantified over number of animals in each condition.

### Cocaine disrupts the correlation between DUSP5 immunoreactivity and synaptic plasticity markers in NAcore MSNs

To determine whether DUSP5 immunoreactivity is associated with structural markers of synaptic plasticity, we performed correlation analyses between DUSP5 signal in dendritic shafts and spines with dendritic spine morphology metrics presented in **Figure 4**. No significant correlation was observed between DUSP5 immunoreactivity—either in dendritic shafts or spines—and spine density (spines per µm) in either saline- or cocaine-treated groups (**Fig. 5A and 5C**). In contrast, in saline-treated animals, DUSP5 immunoreactivity in dendritic spines showed a significant positive correlation with spine head diameter—a key indicator of synaptic strength. This relationship was absent in cocaine-treated rats (**Fig. 5B**). A similar pattern was observed for DUSP5 in dendritic shafts, where a positive correlation with spine head diameter was present in saline controls but abolished in cocaine-treated rodents (**Fig. 5D**). These findings suggest that under baseline conditions, DUSP5 expression is positively associated with spine head size, potentially reflecting a role in maintaining synaptic integrity. Cocaine exposure appears to disrupt this relationship, indicating a cocaine-induced dysregulation of DUSP5-linked signaling pathways. This disruption may contribute to maladaptive synaptic remodeling in NAcore MSNs, which underlies cocaine-seeking behavior.

**Figure 5.**
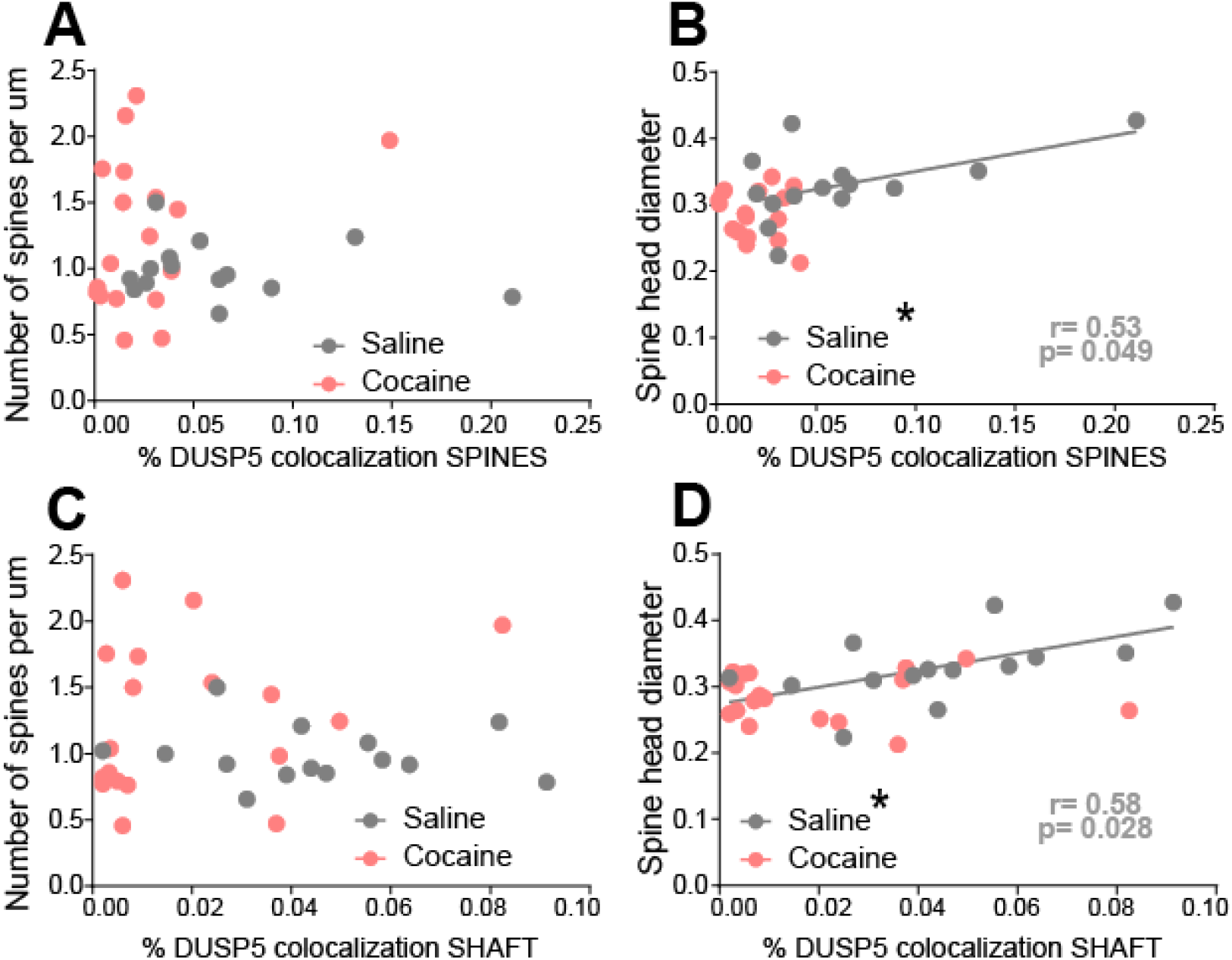
Cocaine-treated rats exhibited a loss of correlation between DUSP5 immunoreactivity in dendritic shafts and spines and synaptic plasticity, as measured by spine head diameter, in NAcore MSNs. Correlations were examined in the same animals analyzed in Figures 2 and 3. **A)** No significant correlation between DUSP5 immunoreactivity in spine heads and spine density (spines/µm) in either saline- or cocaine-treated groups. **B)** A significant positive correlation between DUSP5 immunoreactivity in spine heads and spine head diameter was observed in the saline group, but not in the cocaine group (saline: r = 0.53, p<0.05). **C)** No significant correlation between DUSP5 immunoreactivity in dendritic shafts and spine density in either group. **D)** A significant positive correlation between DUSP5 immunoreactivity in dendritic shafts and spine head diameter was found in the saline group, but not in the cocaine group (saline: r = 0.58, p<0.05).

## DISCUSSION

The present study identifies DUSP5 as a molecular regulator of synaptic plasticity and cocaine-seeking behavior, with effects most prominent in NAcore D1-type MSN. By integrating behavioral pharmacology, high-resolution imaging of dendritic spine morphology, and transcriptomic mining, we provide convergent evidence that deficiencies in DUSP5 regulates relapse-relevant neuroadaptations in a phase- and cell-type-specific manner.

### Dynamic regulation of DUSP5 across the addiction cycle

Data mining of publicly available single-cell RNA sequencing datasets from rats exposed to i.p. cocaine revealed that *DUSP5* mRNA is selectively upregulated in D1-MSN in the NAcore. This transcriptional response was observed 1 hour after non-contingent (i.p.) cocaine administration and was not evident for other DUSP family members or in other cell types, indicating a cell type–specific effect of cocaine on DUSP5 expression. These findings suggest a potential role for DUSP5 in the early molecular adaptations to cocaine exposure and support its involvement in D1-MSN–specific plasticity mechanisms within the NAcore[34]. In contrast, using a well-validated model of cocaine SA and cue-induced reinstatement, we found that DUSP5 protein is significantly downregulated in NAcore MSNs following chronic cocaine exposure, as measured 10 days into abstinence during a reinstatement session. Specifically, DUSP5 immunoreactivity was reduced in both dendritic shafts and spines. This discrepancy between mRNA and protein levels may reflect differences in the timing of measurement (1 hour post-injection vs. 10 days into abstinence), the mode of cocaine exposure (acute, non-contingent vs. chronic, contingent), or post-transcriptional mechanisms affecting DUSP5. Nevertheless, DUSP5 expression was significantly altered by cocaine across different phases of the addiction cycle, highlighting its potential role in cocaine-induced neuroadaptations.

### DUSP5 deficiency correlates with cocaine-seeking behavior

We found that DUSP5 protein levels in dendritic spines and shaft correlated positively with reinstatement behavior, as indexed by active lever presses. Notably, DUSP5 levels did not correlate with total cocaine intake during SA, highlighting its selective involvement in relapse-like behavior rather than cumulative drug exposure. This supports a model in which DUSP5 deficiency contributes to maladaptive plasticity, rather than simply reflecting the pharmacological history of cocaine use.

Furthermore, under control (saline) conditions, DUSP5 levels were positively associated with spine head diameter, a structural correlate of synaptic strength [36, 37]. This relationship was abolished following cocaine exposure, indicating that cocaine disrupts the normal association between DUSP5 and synaptic stability. These findings align with previous work showing that cue-induced cocaine reinstatement is associated with elevated spine head diameter [10, 30, 31, 38] or spine density in NAcore MSNs [39, 40] and reinforce the hypothesis that DUSP5 acts as a homeostatic regulator of excitatory synaptic plasticity.

### Mechanistic insights: DUSP5, p-ERK, and actin remodeling

Our findings reveal a potential mechanistic link between DUSP5 downregulation and cocaine-induced structural plasticity in NAcore MSN. We previously demonstrated that cue-induced reinstatement selectively increases phosphorylated cofilin (p-cofilin) in D1-MSNs, but not in D2-MSNs, supporting the idea that D1-MSN-specific plasticity underlies relapse-like behavior [30]. Cofilin is a key actin-binding protein that regulates the dynamic remodeling of the actin cytoskeleton, a process essential for the formation and stabilization of dendritic spines[41–43]. Its activity is tightly controlled by phosphorylation at serine 3; p-cofilin is inactive and promotes actin filament stabilization and spine head enlargement, both of which are hallmarks of synaptic potentiation [43, 44]. We and others have shown the central role of cofilin in regulating spine morphology and it has emerged as a critical molecular target in studies of drug-induced synaptic plasticity [30, 45, 46]. Thus our data on the selective accumulation of p-cofilin in D1-MSNs aligns with previous reports showing spine head enlargement during cue-induced reinstatement occurs specifically in D1-MSNs [34] and its activation promotes cued drug seeking [9].

DUSP5 may act as a key upstream regulator of the synaptic plasticity process. As a nuclear phosphatase that dephosphorylates ERK1/2, DUSP5 controls ERK signaling intensity and subcellular localization. ERK1/2 can phosphorylate both cofilin and its upstream phosphatase, Slingshot 1 (SSH1); inactivation of SSH1 by ERK sustains p-cofilin accumulation (**Fig. 6**). Thus, loss of dendritic DUSP5 during reinstatement could result in the lost of ERK activity, promoting elevated p-cofilin levels and aberrant spine dynamic, hallmarks of maladaptive plasticity driving relapse. The mined transcriptomic data using non-contigent cocaine injections showed no significant changes in mRNA levels of ERK1/2, LIMK, SSH1, SSH2, or cofilin-1/2, suggesting that post-translational mechanisms, rather than transcriptional regulation, drive the observed structural changes. This positions DUSP5 as a critical post-transcriptional modulator of the ERK–SSH1–cofilin pathway, linking its downregulation directly to synaptic remodeling during relapse. (**Supplemental Fig. 3**)

**Figure 6.**
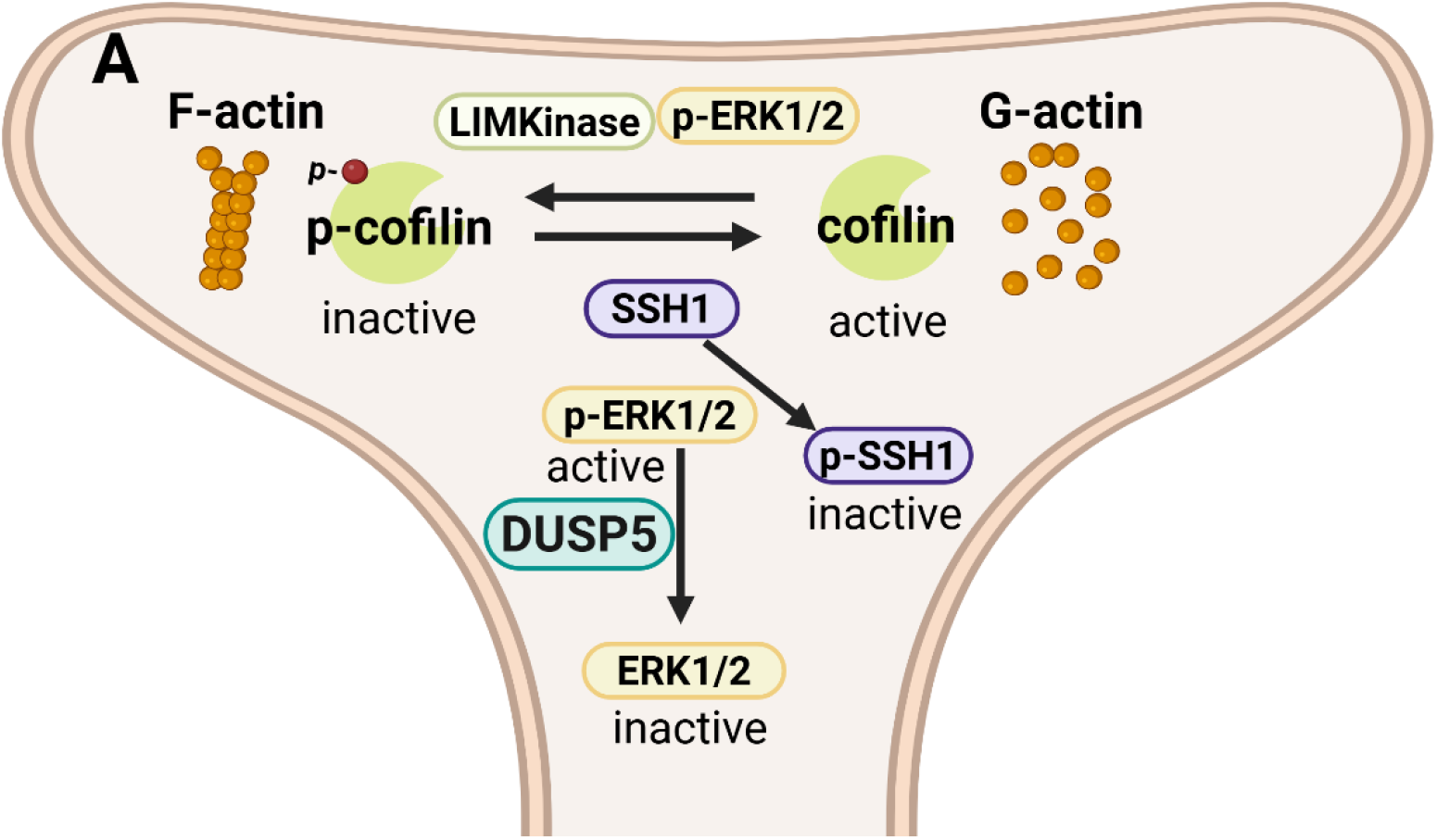
Hypothesized pathway connecting DUSP5 with cofilin, and SSH1 through the interactions with ERK. Cofilin, an actin-binding protein, regulates the dynamic transition between filamentous (F-actin) and globular (G-actin) states. In its dephosphorylated (active) form, cofilin promotes actin depolymerization, whereas phosphorylated cofilin (p-cofilin) is inactive and associated with F-actin stabilization and spine head enlargement and spine number. p-ERK1/2 and LIM kinase phosphorylate cofilin, leading to its inactivation. P-ERK1/2 also phosphorylates and inhibits Slingshot phosphatase 1 (SSH1), the enzyme responsible for cofilin dephosphorylation. DUSP5 negatively regulates this pathway by dephosphorylating p-ERK1/2, thereby limiting its downstream effects on SSH1. Loss of DUSP5 protein, as observed during cue-induced reinstatement, may lead to persistent p-ERK activation, sustained SSH1 inhibition, increased p-cofilin accumulation, and aberrant actin dynamic, a cellular signature of relapse-associated plasticity.

Together, these findings suggest a mechanistic framework in which DUSP5 downregulation disrupts ERK-mediated control of actin dynamics, leading to maladaptive structural plasticity in D1-MSNs during relapse. At present we are collecting experimental evidence to support this hypothesis. This convergence between DUSP5 signaling and actin remodeling pathways underscores a potential molecular pathway by which chronic cocaine exposure alters neuronal function to drive persistent drug-seeking behavior.

### Limitations of the interpretation

While our findings offer new insight into the role of DUSP5 in cocaine-induced synaptic plasticity, some limitations should be acknowledged. First, although we observed a robust downregulation of DUSP5 protein in dendritic compartments during cue-induced reinstatement, our study does not directly demonstrate causality between this reduction and the observed structural or behavioral outcomes. Future experiments using targeted DUSP5 overexpression in D1-MSNs will be essential to establish a functional role in relapse-related plasticity.

Second, we rely on correlation analyses to link DUSP5 expression with spine morphology and cocaine-seeking behavior. While the observed dissociation between DUSP5 levels and cumulative cocaine intake supports a specific role in relapse, correlational data cannot rule out the influence of other upstream or parallel mechanisms, including additional phosphatases or kinases that act on the ERK–SSH1–cofilin pathway.

Third, our model integrates transcriptomic and protein-level data collected at distinct time points and under different cocaine exposure paradigms. Specifically, the snRNA-seq dataset reflects acute, non-contingent cocaine exposure measured 1 hour post-injection, while our immunohistochemical analyses were performed after chronic, self-administered cocaine exposure and 10 days of abstinence, followed by reinstatement. This temporal and methodological mismatch may underlie the observed divergence between DUSP5 mRNA upregulation and dendritic protein downregulation, and underscores the need for time-resolved, cell-type– specific measurements of DUSP5 at both the transcript and protein levels throughout the addiction cycle.

Finally, while our data suggest that post-translational regulation via ERK–SSH1–cofilin signaling is a likely mechanism linking DUSP5 downregulation to altered spine dynamics, we did not directly measure ERK or p-cofilin levels in the same neurons analyzed for DUSP5. Future studies should integrate multiplexed imaging or biochemical approaches to dissect the spatial and temporal coordination of these molecular changes within D1-MSNs. These limitations highlight the need for further mechanistic work to validate DUSP5 as a key modulator of cocaine-induced neuroplasticity and to determine its therapeutic potential in relapse prevention.

## CONCLUSION

Using a well-validated model of cocaine SA and cue-induced reinstatement, we observed a striking downregulation of DUSP5 protein in dendritic spines and shafts of NAcore MSNs, and single-nucleus transcriptomic mined data showed selective upregulation of *DUSP5* mRNA in D1-MSNs following acute cocaine exposure. This transcription–protein dissociation suggests complex, phase-specific regulation of DUSP5 across the addiction cycle. Importantly, DUSP5 protein levels did not correlate with total cocaine intake, but instead showed a strong association with cue-induced reinstatement behavior, reinforcing its role in relapse vulnerability rather than in cumulative drug exposure. Furthermore, cocaine disrupted the normal correlation between DUSP5 and spine head diameter and spine density, a structural hallmark of synaptic potentiation, implicating DUSP5 loss in maladaptive plasticity mechanisms. Overall, this work highlights DUSP5 as a promising molecular target for interventions aimed at disrupting the synaptic plasticity that underlies cocaine relapse.

## Author Contributions

D.C.C, R.R and C.G.K. conceptualized the paper. The analysis was conducted by J.P.T. and M.M. The manuscript was prepared, reviewed, and edited by D.C.C, R.R and C.G.K. All authors have read and agreed to the published version of the manuscript.

## Institutional Review Board Statement

Not applicable.

## Data Availability Statement

The original contributions presented in this study are included in the article material. Further inquiries can be directed to the corresponding authors.

## Conflicts of Interest

The authors declare no conflicts of interest. RR. is a co-founder of CIAN.

### Statistic analysis extracted form [32]

Cocaine administration increased Dusp5 mRNA expression in DDR1-expressing MSN. **(acute cocaine: p-value <0.00001, log2FoldChange= 2.778; repeated cocaine: p-value= 0.509, log2FoldChange= 0.416)**. Cocaine did not significantly change the mRNA levels of Dusp6 in DDR1-msns-1 (repeated cocaine: p-value= 0.747, log2FoldChange= −0.303; acute cocaine: p-value =0.001 log2FoldChange= 2.837). Cocaine did not change the mRNA levels of Dusp7 in DDR1-msns-1 (repeated cocaine: p-value= 0.998, log2FoldChange= 0.006; acute cocaine: p-value =0.093 log2FoldChange= −0.908). Cocaine did not change the mRNA levels of Dusp14 in DDR1-msns-1 (repeated cocaine: p-value= 0.495, log2FoldChange= 0.181; acute cocaine: p-value =0.018 log2FoldChange= 0.758). Cocaine did not change the mRNA levels of Dusp15 in DDR1-msns-1 (repeated cocaine: p-value= 0.766, log2FoldChange= 0.123; acute cocaine: p-value =0.819 log2FoldChange= −0.115). Cocaine did not change the mRNA levels of Limk1 in DDR1-msns-1 (repeated cocaine: p-value= 0.741, log2FoldChange= 0.179; acute cocaine: p-value =0.742 log2FoldChange= −0.162). Cocaine did not change the mRNA levels of Limk2 in DDR1-msns-1 (repeated cocaine: p-value= 0.572, log2FoldChange= −0.125; acute cocaine: p-value =0.278 log2FoldChange= −0.271). Cocaine did not change the mRNA levels of Mapk1 in DDR1-msns-1 (repeated cocaine: p-value= 0.609, log2FoldChange= −0.059; acute cocaine: p-value =0.741, log2FoldChange= −0.044).

## REFERENCES

1. Friedman, J. and C.L. Shover, Charting the fourth wave: Geographic, temporal, race/ethnicity and demographic trends in polysubstance fentanyl overdose deaths in the United States, 2010-2021. Addiction, 2023. 118(12): p. 2477–2485.

2. Ciccarone, D., The rise of illicit fentanyls, stimulants and the fourth wave of the opioid overdose crisis. Curr Opin Psychiatry, 2021. 34(4): p. 344–350.

3. Dasgupta, N., L. Beletsky, and D. Ciccarone, Opioid Crisis: No Easy Fix to Its Social and Economic Determinants. Am J Public Health, 2018. 108(2): p. 182–186.

4. Volkow, N.D. and C. Blanco, Substance use disorders: a comprehensive update of classification, epidemiology, neurobiology, clinical aspects, treatment and prevention. World Psychiatry, 2023. 22(2): p. 203–229.

5. Garcia-Keller, C., et al., Glutamatergic mechanisms of comorbidity between acute stress and cocaine self-administration. Mol Psychiatry, 2016. 21(8): p. 1063–9.

6. Kelley, A.E., Ventral striatal control of appetitive motivation: role in ingestive behavior and reward-related learning. Neurosci Biobehav Rev, 2004. 27(8): p. 765–76.

7. Scofield, M.D., et al., The Nucleus Accumbens: Mechanisms of Addiction across Drug Classes Reflect the Importance of Glutamate Homeostasis. Pharmacol Rev, 2016. 68(3): p. 816–71.

8. Gerfen, C.R. and D.J. Surmeier, Modulation of striatal projection systems by dopamine. Annu Rev Neurosci, 2011. 34: p. 441–66.

9. Pardo-Garcia, T.R., et al., Ventral Pallidum Is the Primary Target for Accumbens D1 Projections Driving Cocaine Seeking. J Neurosci, 2019. 39(11): p. 2041–2051.

10. Gipson, C.D., et al., Relapse induced by cues predicting cocaine depends on rapid, transient synaptic potentiation. Neuron, 2013. 77(5): p. 867–72.

11. Park, M., et al., Plasticity-induced growth of dendritic spines by exocytic trafficking from recycling endosomes. Neuron, 2006. 52(5): p. 817–30.

12. Oh, W.C., T.C. Hill, and K. Zito, Synapse-specific and size-dependent mechanisms of spine structural plasticity accompanying synaptic weakening. Proc Natl Acad Sci U S A, 2013. 110(4): p. E305–12.

13. Sugiura, R., R. Satoh, and T. Takasaki, ERK: A Double-Edged Sword in Cancer. ERK-Dependent Apoptosis as a Potential Therapeutic Strategy for Cancer. Cells, 2021. 10(10).

14. Yap, J.L., et al., Small-molecule inhibitors of the ERK signaling pathway: Towards novel anticancer therapeutics. ChemMedChem, 2011. 6(1): p. 38–48.

15. Imhoff, A., et al., Structural and kinetic characterization of DUSP5 with a Di-phosphorylated tripeptide substrate from the ERK activation loop. Front Chem Biol, 2024. 3.

16. Kuntz-Melcavage, K.L., et al., Gene expression changes following extinction testing in a heroin behavioral incubation model. BMC Neurosci, 2009. 10: p. 95.

17. Seleman, M., et al., Impact of P-glycoprotein at the blood-brain barrier on the uptake of heroin and its main metabolites: behavioral effects and consequences on the transcriptional responses and reinforcing properties. Psychopharmacology (Berl), 2014. 231(16): p. 3139–49.

18. Ujike, H., et al., Gene expression related to synaptogenesis, neuritogenesis, and MAP kinase in behavioral sensitization to psychostimulants. Ann N Y Acad Sci, 2002. 965: p. 55–67.

19. Takaki, M., et al., Two kinds of mitogen-activated protein kinase phosphatases, MKP-1 and MKP-3, are differentially activated by acute and chronic methamphetamine treatment in the rat brain. J Neurochem, 2001. 79(3): p. 679–88.

20. Qiao, X., et al., Dual-specificity phosphatase 15 (DUSP15) in the nucleus accumbens is a novel negative regulator of morphine-associated contextual memory. Addict Biol, 2021. 26(1): p. e12884.

21. Nielsen, D.A., et al., Genome-wide association study identifies genes that may contribute to risk for developing heroin addiction. Psychiatr Genet, 2010. 20(5): p. 207–14.

22. Kutty, R.G., et al., Dual Specificity Phosphatase 5-Substrate Interaction: A Mechanistic Perspective. Comprehensive Physiology, 2017. 7(4): p. 1449–1461.

23. Talipov, M.R., et al., Critical Role of the Secondary Binding Pocket in Modulating the Enzymatic Activity of DUSP5 toward Phosphorylated ERKs. Biochemistry, 2016. 55(44): p. 6187–6195.

24. Imhoff, A., et al., Structural and kinetic characterization of DUSP5 with a Di-phosphorylated tripeptide substrate from the ERK activation loop. Frontiers in Chemical Biology, 2024. In press.

25. Piras, I.S., et al., Whole transcriptome profiling of the human hippocampus suggests an involvement of the KIBRA rs17070145 polymorphism in differential activation of the MAPK signaling pathway. Hippocampus, 2017. 27(7): p. 784–793.

26. https://www.proteinatlas.org/ENSG00000138166-DUSP5/brain/hippocampal%2Bformation?utm.

27. Garcia-Keller, C., et al., Extracellular Matrix Signaling Through β3 Integrin Mediates Cocaine Cue– Induced Transient Synaptic Plasticity and Relapse. Biological Psychiatry, 2019. 86(5): p. 377–387.

28. Garcia-Keller, C., et al., Relapse-Associated Transient Synaptic Potentiation Requires Integrin-Mediated Activation of Focal Adhesion Kinase and Cofilin in D1-Expressing Neurons. The Journal of Neuroscience, 2020. 40(44): p. 8463–8477.

29. Akiki, R.M., et al., A long noncoding eRNA forms R-loops to shape emotional experience-induced behavioral adaptation. Science, 2024. 386(6727): p. 1282–1289.

30. Garcia-Keller, C., et al., Relapse-Associated Transient Synaptic Potentiation Requires Integrin-Mediated Activation of Focal Adhesion Kinase and Cofilin in D1-Expressing Neurons. J Neurosci, 2020. 40(44): p. 8463–8477.

31. Garcia-Keller, C., et al., Extracellular Matrix Signaling Through beta3 Integrin Mediates Cocaine Cue-Induced Transient Synaptic Plasticity and Relapse. Biol Psychiatry, 2019. 86(5): p. 377–387.

32. Phillips, R.A., 3rd, et al., Distinct subpopulations of D1 medium spiny neurons exhibit unique transcriptional responsiveness to cocaine. Mol Cell Neurosci, 2023. 125: p. 103849.

33. Spencer, S., et al., Cocaine Use Reverses Striatal Plasticity Produced During Cocaine Seeking. Biol Psychiatry, 2017. 81(7): p. 616–624.

34. Bobadilla, A.C., et al., Corticostriatal plasticity, neuronal ensembles, and regulation of drug-seeking behavior. Prog Brain Res, 2017. 235: p. 93–112.

35. Gipson, C.D., et al., Reinstatement of nicotine seeking is mediated by glutamatergic plasticity. Proceedings of the National Academy of Sciences of the United States of America, 2013. 110(22): p. 9124–9.

36. Borczyk, M., et al., Neuronal plasticity affects correlation between the size of dendritic spine and its postsynaptic density. Sci Rep, 2019. 9(1): p. 1693.

37. Hotulainen, P. and C.C. Hoogenraad, Actin in dendritic spines: connecting dynamics to function. J Cell Biol, 2010. 189(4): p. 619–29.

38. Shen, H.W., et al., Prelimbic cortex and ventral tegmental area modulate synaptic plasticity differentially in nucleus accumbens during cocaine-reinstated drug seeking. Neuropsychopharmacology, 2014. 39(5): p. 1169–77.

39. Robinson, T.E. and B. Kolb, Structural plasticity associated with exposure to drugs of abuse. Neuropharmacology, 2004. 47 Suppl 1: p. 33–46.

40. Russo, S.J., et al., The addicted synapse: mechanisms of synaptic and structural plasticity in nucleus accumbens. Trends Neurosci, 2010. 33(6): p. 267–76.

41. Ono, S., Regulation of actin filament dynamics by actin depolymerizing factor/cofilin and actin-interacting protein 1: new blades for twisted filaments. Biochemistry, 2003. 42(46): p. 13363–70.

42. Paciello, F., et al., Role of LIMK1-cofilin-actin axis in dendritic spine dynamics in Alzheimer’s disease. Cell Death Dis, 2025. 16(1): p. 431.

43. Rust, M.B., ADF/cofilin: a crucial regulator of synapse physiology and behavior. Cell Mol Life Sci, 2015. 72(18): p. 3521–9.

44. Noguchi, J., et al., State-dependent diffusion of actin-depolymerizing factor/cofilin underlies the enlargement and shrinkage of dendritic spines. Sci Rep, 2016. 6: p. 32897.

45. Kruyer, A., et al., Post-translational S-glutathionylation of cofilin increases actin cycling during cocaine seeking. PLoS One, 2019. 14(9): p. e0223037.

46. Toda, S., et al., Cocaine increases actin cycling: effects in the reinstatement model of drug seeking. J Neurosci, 2006. 26(5): p. 1579–87.

