## Supplemental Materials for "DUSP5 Downregulation in Nucleus Accumbens Core Correlates with Synaptic Plasticity and Cue-Induced Cocaine Reinstatement"

**Supplemental files**

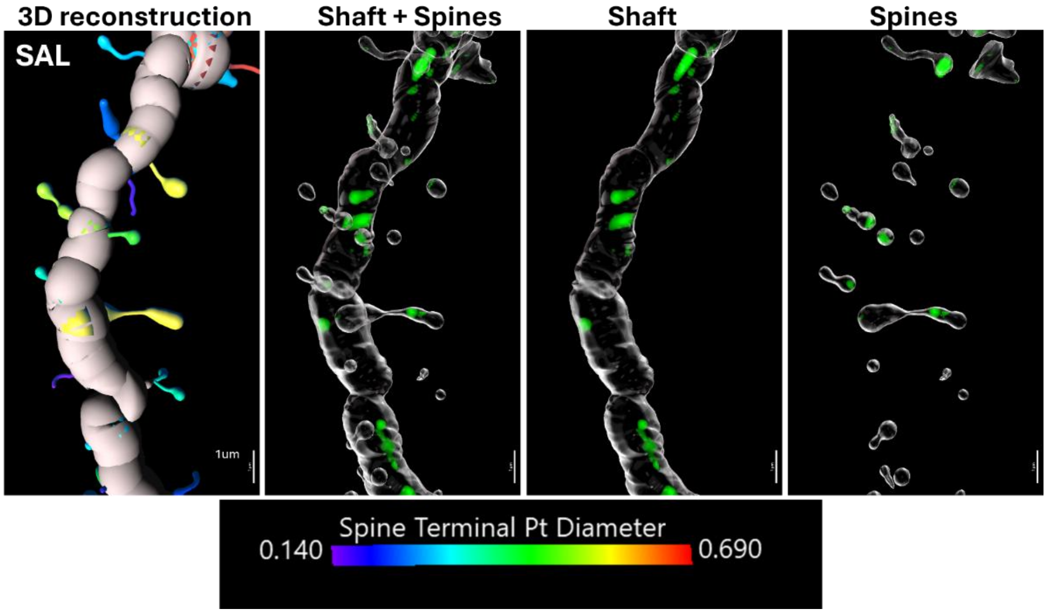


**Supplemental Figure 1. Quantifying dendritic DUSP5 protein immunoreactivity in labeled MSNs from a saline-treated control animal.** This figure presents the rendering and quantification of a confocal image showing a virus-labeled dendritic segment from a medium spiny neuron (MSN). **Panel 1** illustrates the dendritic structure filled using the filament module in the IMARIS software, with spine heads color-coded based on their diameters. **Panel 2** displays a 3D-flattened, masked representation of the dendritic shaft and spines (in transparent white), overlaid with the surrounding DUSP5 signal (green). **Panel 3** isolates the 3D-flattened, masked dendritic shaft with the adjacent DUSP5 signal, while **Panel 4** focuses on the 3D-flattened, masked dendritic spines and their surrounding DUSP5 signal.

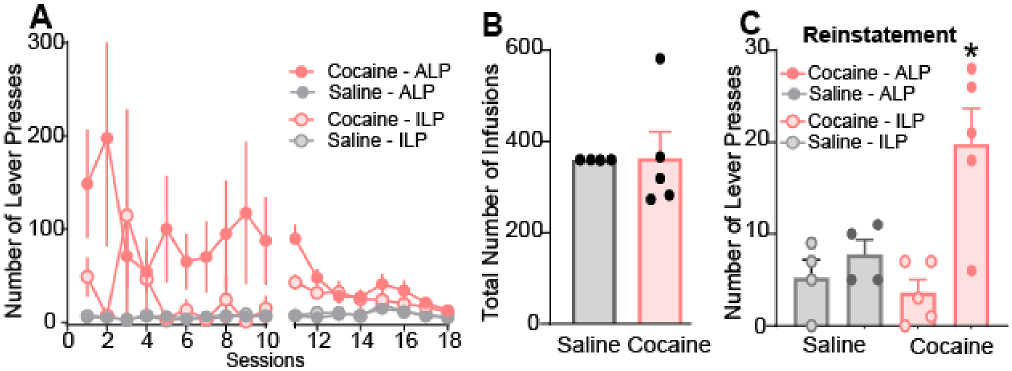


**Supplemental Figure 2. A)** Time course of active and inactive lever pressing of SA and extinction in saline and cocaine- and saline-treated rats used for DUSP5 immunoreactivity. [active lever presses: two-way ANOVA repeated measures over time, time $F_{(2.29,16.06)}$= 1.08, $p$=0.37; treatment $F_{(1,7)}$ = 6.65, $p$ = 0.036; interaction $F_{(17,119)}$ = 1.162, $p$ = 0.30; inactive lever presses: two-way ANOVA repeated measures over time, time $F_{(1.064,7.45)}$= 0.64, $p$=0.64; treatment $F_{(1,7)}$ = 1.86, $p$ = 0.215; interaction $F_{(17,119)}$ = 0.678, p = 0.81]. **B)** Total number of infusions for saline and cocaine-treated rats. **C)** Number of active and inactive lever presses during cue-induced reinstatement. [ two-way ANOVA, treatment $F_{(1, 14)}$= 4.011, $p$ = 0.065; active vs inactive $F_{(1,14)}$ = 12.97, $p$ = 0.003; interaction $F_{(1,14)}$ = 6.96, $p$ = 195].

Data are shown as mean ± SEM. *N* correspond to 5 cocaine animals and 4 saline animals. ALP= active lever presses, ILP= inactive lever presses.

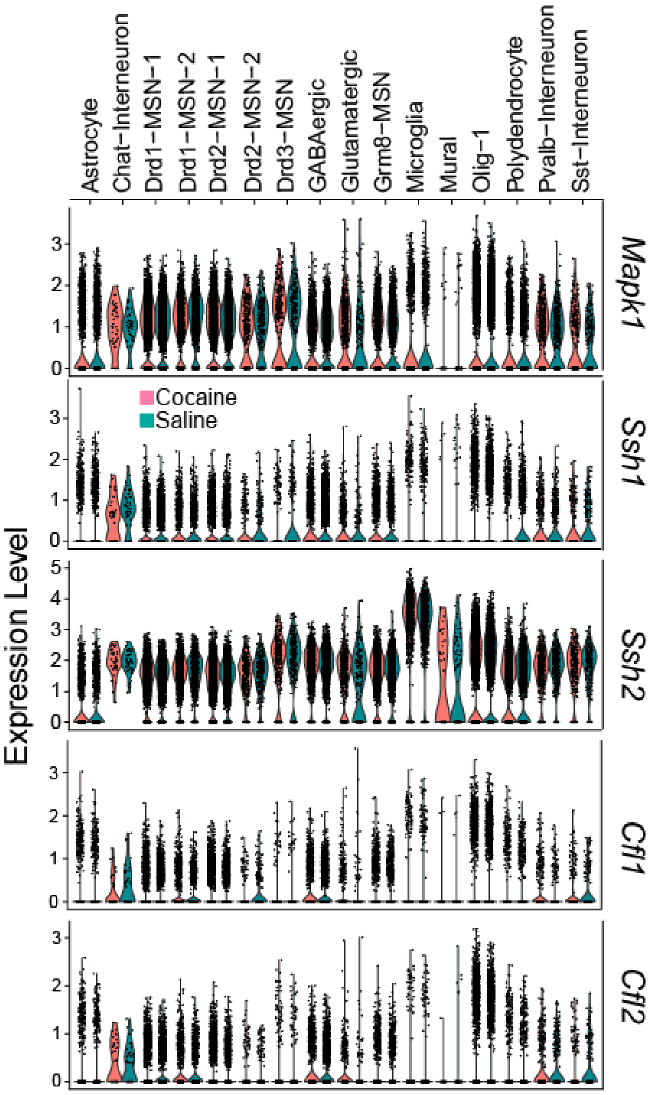


**Supplemental Figure 3. Single-cell RNA sequencing analysis shows stable expression of actin-regulatory and ERK pathway genes across cell types following acute cocaine exposure.** Violin plots display gene expression levels of CFL1, CFL2 (*cofilin isoforms*), SSH1, SSH2 (*Slingshot phosphatases*), and MAPK1 (*ERK2*) across major brain cell types in the nucleus accumbens (NAc), comparing rats treated with acute intraperitoneal cocaine (pink) or saline (teal). Data were obtained from a publicly available single-cell RNA-sequencing dataset [32] collected 1 hour post-injection. Cell types include D1-MSNs, D2-MSNs, GABAergic interneurons, astrocytes, oligodendrocytes, mural cells, endothelial cells, and others. Across all genes and cell types examined, no significant differences in mRNA expression were observed between saline and cocaine groups, including in D1-MSNs, the cell type implicated in cocaine-induced plasticity. These findings suggest that the ERK–SSH–cofilin signaling pathway is not transcriptionally altered at this time point, pointing instead to post-transcriptional or post-translational regulation (e.g., phosphorylation), consistent with DUSP5's role as a phosphatase acting on ERK1/2.
